# Coumarin as a general switching auxiliary to prepare photochromic and spontaneously blinking fluorophores

**DOI:** 10.1101/2024.05.12.593749

**Authors:** Fadi M. Jradi, Brian P. English, Timothy A. Brown, Jesse Aaron, Satya Khuon, James A. Galbraith, Catherine G. Galbraith, Luke D. Lavis

## Abstract

Single-molecule localization microscopy (SMLM) uses activatable or switchable fluorophores to create non-diffraction limited maps of molecular location in biological samples. Despite the utility of this imaging technique, the portfolio of appropriate labels for SMLM remains limited. Here, we describe a general strategy for the construction of “glitter bomb” labels by simply combining rhodamine and coumarin dyes though an amide bond. Condensation of the *ortho*-carboxyl group on the pendant phenyl ring of rhodamine dyes with a 7-aminocoumarin yields photochromic or spontaneously blinking fluorophores depending on the parent rhodamine structure. We apply this strategy to prepare labels useful super-resolution experiments in fixed cells using different attachment techniques. This general glitter bomb strategy should lead to improved labels for SMLM, ultimately enabling the creation of detailed molecular maps in biological samples.

## INTRODUCTION

Fluorescence imaging enables interrogation of cellular structures in cells with molecular specificity.^1-3^ The advent of super-resolution techniques, such as single-molecule localization microscopy (SMLM),^4-6^ has pushed the resolution of optical imaging near that of electron microscopy. Fluorescence imaging relies on detecting the light emitted from fluorescent proteins (FPs)^7^ or small-molecule fluorophores^8^ bound to the biological target of interest.^9^ For SMLM, the emission of individual fluorophores must also be separable in time, which can be achieved though photochemical activation, lig- and binding, enzymatic catalysis, or transient protonation. Although a variety of photoactivatable and photoswitchable FPs have been described, small-molecule fluorophores are often preferred for SMLM due to their chromatic modularity^10^ and large photon outputs.^11, 12^

To date, the majority of SMLM experiments utilize *direct* stochastic optical reconstruction microscopy (*d*STORM), where conventional cyanine dyes are switched on and off in a photoredox process under intense illumination and strong reducing buffers.^13-15^ *d*STORM has led to new biological discoveries,^16, 17^ but requires specific chemical conditions to elicit photoswitching. This limitation has prompted the development of fluorophores that cycle between on- and off-states without the need for reducing conditions.^18^ An attractive scaffold for such fluorophores is the rhodamine dye class where the *ortho*-carboxylate nucleofuge/nucleophile leaves/attacks the xanthene system in a reversible manner (**Figure 1a**). This results in an equilibrium between a nonfluorescent lactone form and fluorescent zwitterion. The lactone–zwitterion equilibrium constant (*K*_L–Z_), is dependent on the rhodamine structure and a key predictor of the performance of rhodamines in biological systems.^19^

**Figure 1.**
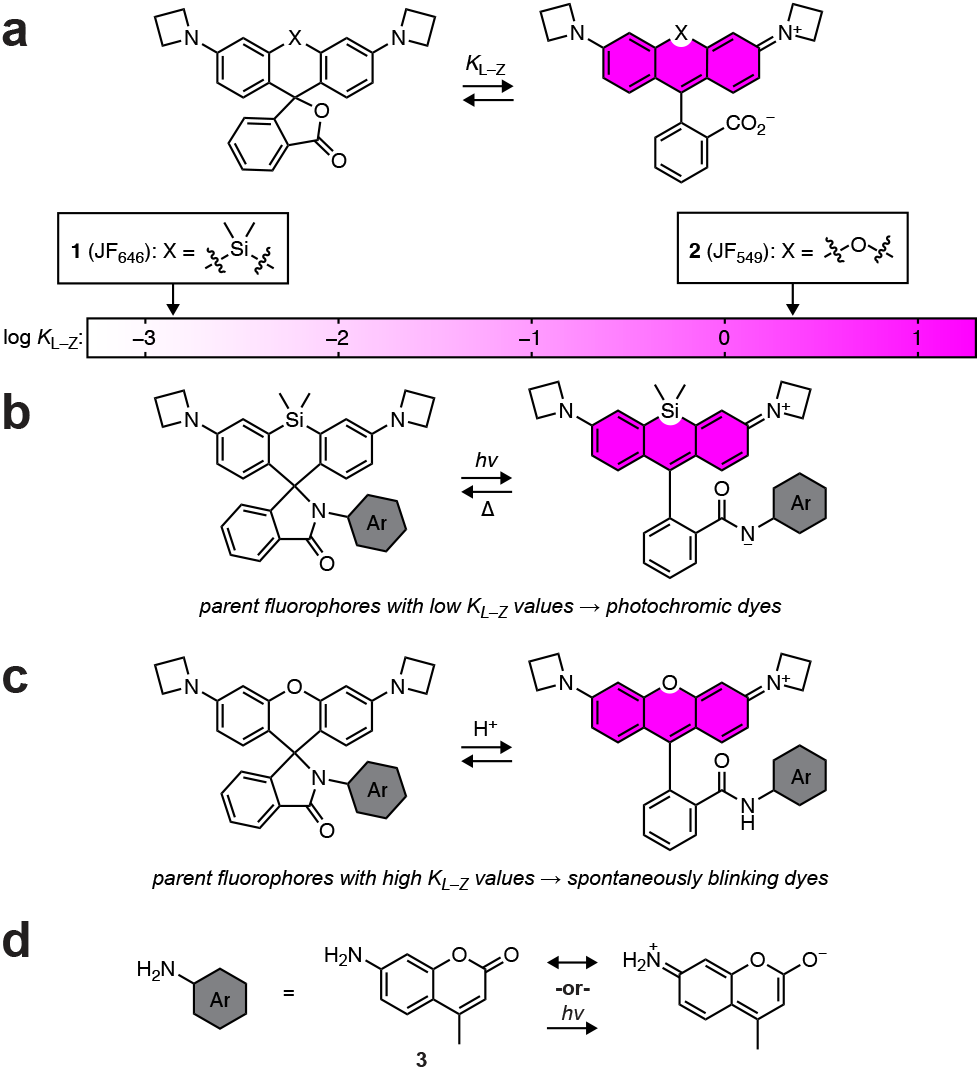
Coumarin as a universal switch for rhodamine dyes. (**a**) The lactone–zwitterion equilibrium of rhodamine dyes and the chemical structures of JF_646_ (**1**), which exhibits a low lactone–zwitterion equilibrium constant (*K*_L–Z_) and JF_549_ (**2**) that shows a high *K*_L–Z_. (**b**) Photochromic dye based on JF_646_ conjugated to an appropriate aryl amine auxiliary (Ar). (**c**) Spontaneously blinking fluorophore based on JF_549_ aminated with an appropriate ArNH_2_ switching auxiliary group. (**d**) 7-Amino-4-methylcoumarin as a general ArNH_2_ group due to charge transfer in both the ground state and the excited state.

For rhodamines containing a Si(CH_3_)_2_ bridging substituent, such as Janelia Fluor 646 (JF_646_, **1, Figure 1a**),^12, 20^ the molecule preferentially adopts the nonfluorescent zwitterionic form in aqueous solution, resulting a low *K*_L–Z_ value. The relatively high electrophilicity of the dibenzosilinylium group can be exploited to create spontaneously blinking dyes by reducing the *ortho*-carboxylate to a hydroxymethyl (HM) group.^21^ These dyes undergo transient protonation/deprotonation with p*K*_a_ near 6, resulting in a dye that spontaneously blinks at physiological pH. In contrast, oxygen-containing, xanthene-based rhodamines like Janelia Fluor 549 (JF_549_, **2**) favor the fluorescent zwitterionic form yielding a higher *K*_L–Z_ value. The HM strategy can also be applied to classic rhodamines but the lower electrophilicity of the xanthylium moiety requires additional modifications, such as acylation of the amino auxochromes^21^ or fluorination^22-24^ to create a dye an appropriate duty cycle for super-resolution imaging. Overall, the highly variable properties of rhodamines complicate the development of labels for SMLM since each dye requires specific functionality to tune switching behavior.

We sought a general method for creating rhodamine-based labels for SMLM. We envisioned condensing the *ortho*-carboxyl of rhodamine dyes with an aryl amine switching auxiliary that could serve two functions. For rhodamines that preferentially adopt the lactone form, such as Si-rhodamine **1**, amidation of the carboxyl group to install an appropriate *N*-aryl group could give a photochromic derivative that can be transiently activated with violet light and then thermally relax to the nonfluorescent form (**Figure 1b**). This strategy has been applied to standard rhodamine dyes^25, 26^ but has not been applied to Si-rhodamines. For dyes that exhibit high *K*_L–Z_ values, including many oxygen-containing rhodamines like JF_549_ (**2**), amidation with an appropriate aryl amine moiety could shift the equilibrium to the lactam, but yield a spontaneously blinking dye where transient protonation switches the dye between nonfluorescent and fluorescent forms (**Figure 1c**). Spontaneously blinking fluorophores have been prepared by amidation of the *ortho*-carboxyl group on tetramethylrhodamine with an electron-deficient alkyl amine.^27^ Si-rhodamines where the pendant phenyl ring is replaced with a thiophene to increase *K*_L–Z_ can be amidated with arylamines to create spontaneously blinking fluorophores.^28^

To choose an appropriate *N*-aryl group we considered both photochromic rhodamines and spontaneously blinking fluorophores from a physical organic chemistry perspective. We recognized that photochromism is dictated by the nucleophilicity of the aryl amide in the excited-state whereas spontaneously blinking behavior is governed by the nucleophilicity of the *N*-aryl amide in the ground-state. We were curious if modulation of both ground-state and excited-state nucleophilicity could be achieved with a single chemical motif. We considered coumarins, which exhibit enormous chemical diversity^29, 30^ and are used as pharmacological agents,^31^ photolabile groups,^32^ photosenstizers,^33^ fluorescent indicators^34^ as well as fluo-rescent and fluorogenic probes.^35^ This broad use stems from their straightforward syntheses,^36-38^ which allow modulation of both chemical and optoelectronic properties. We hypothesized that the classic 7-aminocoumarin system could serve as a universal switch for both photochromic and spontaneously blinking dyes.

This idea is predicated on the conjugated donor–acceptor (D–A) coumarin chromophore system. The resonance structures of the prototypical 7-amino-4-methylcoumarin (**3**; **Figure 1d**) indicates low nucleophilicity of the aniline nitrogen in both the excited state and ground state. Here, report the rational design of new SMLM dyes synthesized by simply amidating known rhodamine dyes with a 7-aminocoumarin. These include a photochromic derivative of the far-red Si-rhodamine, Janelia Fluor 646 (JF_646_) and spontaneously blinking derivatives of Janelia Fluor 549 (JF_549_) and Alexa Fluor 594 (AF_594_). We demonstrate the utility of these new “glitter bomb” (GB) dyes using different targeting strategies^9^ to achieve SMLM in fixed-cell imaging experiments.

## RESULTS AND DISCUSSION

### The *K*_L–Z_ of rhodamine dyes predicts the p*K*_a_ of coumarin amide derivatives

To test our hypothesis that coumarin could serve a dual role depending on the rhodamine partner, we sought to synthesized a panel of rhodamine amides using Coumarin 151 (**4**) as the *N*-aryl auxiliary. Coumarin 151 is an established laser dye and the CF_3_ group improves photostability, elicits a modest bathochromic shift, and reinforces the D–A nature of the chromophore.^39, 40^ With parent dyes that exhibit low *K*_L–Z_ values, such as JF_646_ (**1, Figure 1a**), we expected the corresponding rhodamine–coumarin amide adducts to be difficult to protonate in aqueous solution and show low p*K*_a_ values. As explained above, such compounds could serve as photochromic fluorophores as excitation of the coumarin moiety could decrease the nucleophilicity of the coumarin nitrogen, yielding a transient fluorescent species (**Figure 1b**). The coumarin adducts of dyes with higher *K*_L–Z_ values (*e*.*g*., JF_549_, **2**) should show correspondingly higher p*K*_a_ values. In this case, such compounds could be transiently protonated near neutral pH, yielding spontaneously blinking fluorophores.

To examine the relationship between *K*_L–Z_ of the parent dye and p*K*_a_ of a rhodamine–coumarin adduct, we first tested conditions to condense coumarin **4** to various rhodamines, finding that either (COCl)_2_ or HATU could be used to form the desired amide bond depending on the rhodamine starting material. For example, treatment of JF_646_ (**1**) with (COCl)_2_ followed by addition of **4** yielded lactam **5** (**Figure 2a**). We then conjugated **4** to six other rhodamine dyes with *K*_L–Z_ values that range between 0.068 and 9.32 in 1:1 dioxane:H_2_O mixtures generating rhodamine–coumarin lactams **6**–**11** (**Figure 2b**). We then measured the p*K*_a_ of these rhodamine–coumarin compounds, discovering a clear correlation between the *K*_L–Z_ values of the parent dye and the p*K*_a_ of the lactam (**Figure 2c,d**). Rhodamine–coumarin conjugates based on parent dyes with log *K*_L–Z_ < 0 remained in the colorless lactam form, even at pH = 2. These compounds included the coumarin adducts of Si-rhodamine JF_646_ (**5**),^12^ 3,3-difluorazetidinyl coumarin JF_525_ (**6**),^20^ and the fluorinated Si-rhodamine JF_669_ (**7**).^19^ Compounds **8**–**11** are based on rhodamine dyes with log *K*_L–Z_ > 0. Rhodamine–coumarin **8**, which is derived from JF_549_ (**1**; *K*_L–Z_ = 3.47), exhibited a p*K*_a_ = 2.9. Compound **9**, based on Rhodamine 110 (Rh_110_; *K*_L–Z_ = 4.11), showed a higher p*K*_a_ = 3.5. We also investigated more substituted rhodamine dyes such as the sulfonated Alexa Fluor 594 (AF_594_; *K*_L–Z_ = 5.85); the lactam derivative **10** shows p*K*_a_ = 3.7. Finally, lactam **11**, derived from rhodamine 101 (Rh_101_; *K*_L–Z_ = 9.32) showed a higher p*K*_a_ = 4.1. Compounds **7**– **11** show a linear relationship between the p*K*_a_ of the dye amide and the *K*_L–Z_ of the parent fluorophore (**Figure 2c**).

**Figure 2.**
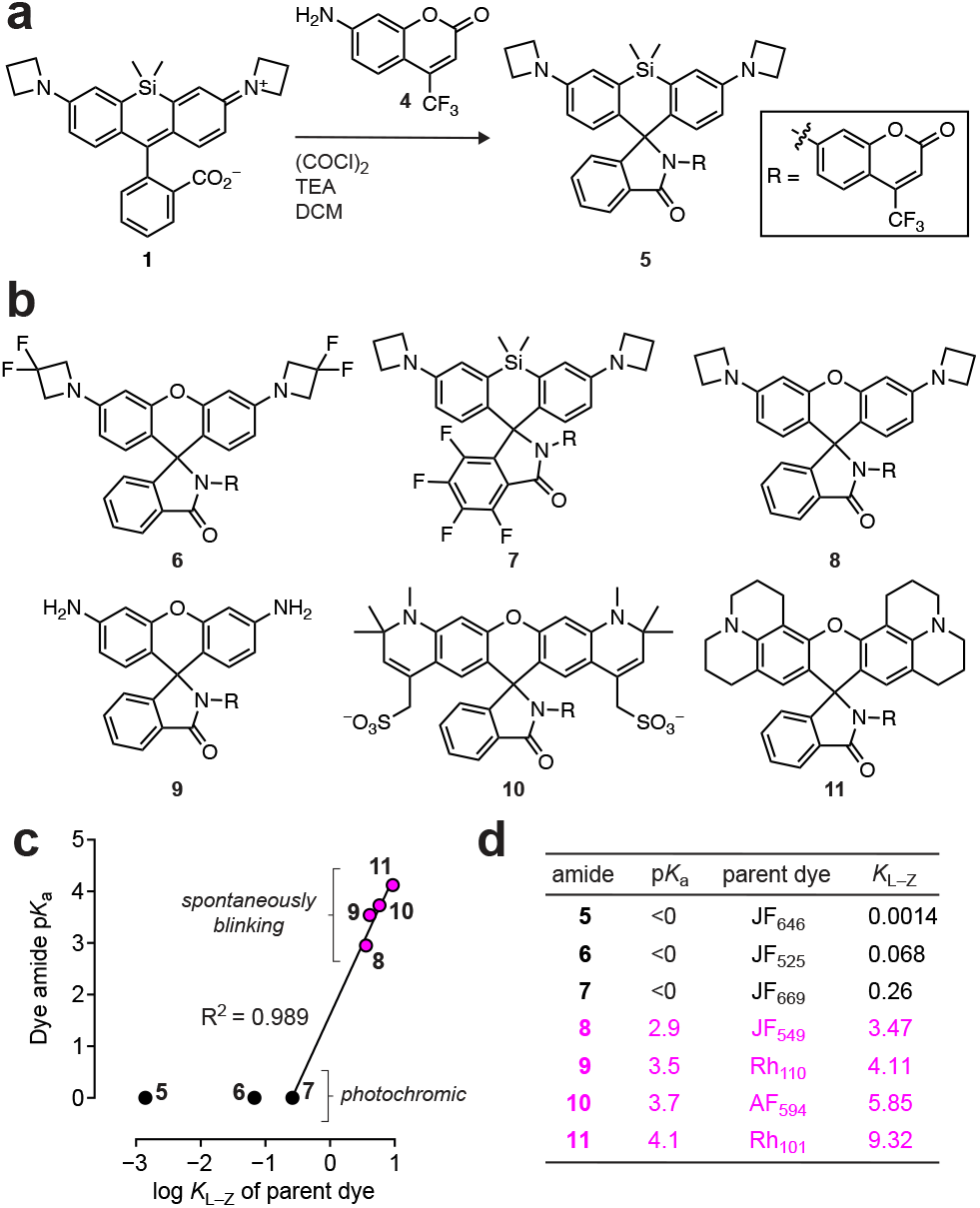
The *K*_L–Z_ of rhodamine dyes predicts the p*K*_a_ of coumarin– rhodamine adducts. (**a**) Condensation of JF_646_ (**1**) and Coumarin 151 (**4**) to prepare GB_646_ (**5**). (**b**) Chemical structures of rhodamine–coumarin amides **6**–**11**. (**c**) Plot of rhodamine–coumarin amide p*K*_a_ *vs*. log *K*_L–Z_ of parent rhodamine dye indicating potential photochromic dyes (black circles) and spontaneously blinking fluorophores (magenta circles). (**d**) Table of properties for compounds **5**–**11**.

### A photochromic Si-rhodamine for SMLM based on JF_646_

We initially focused on potential photochromic derivatives and considered the established photochromic rhodamine **12**,^25^ which consists of a Rhodamine B conjugated to a phthalimide derivative (**Figure 3a**). We compared this to our Si-rhodamine–coumarin compound **5**. In our model of rhodamine photochromism, the phthalimide or coumarin D–A systems undergo photoinduced intramolecular charge transfer (ICT) upon excitation to the first singlet excited state,^41^ enhancing their dipole moment.^42^ This decreases the nucleophilicity of the amide nitrogen,^43-45^ breaking the lactam bond and generating a transient rhodamine species that is fluorescent. This short-lived intermediate then relaxes thermally to the colorless and nonfluorescent lactam (**Figure 1b**).^46^

**Figure 3.**
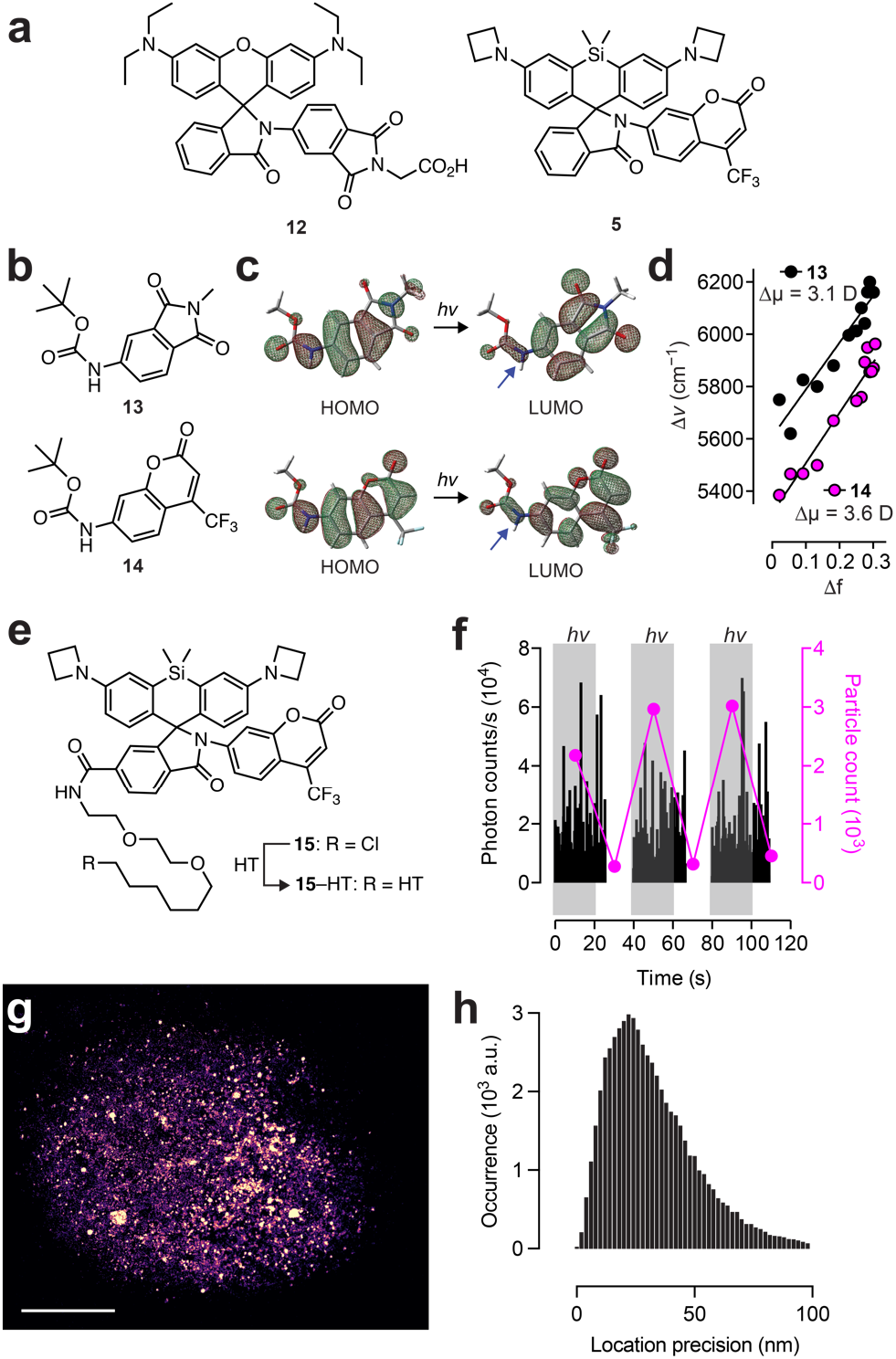
Coumarin is an appropriate switching auxiliary for photochromic rhodamine dyes. (**a**) Chemical structures of rhodamine B– phthalimide **12** and GB_646_ (**5**). (**b**) Chemical structures of model amides **13** and **14**. (**c**) DFT calculated HOMO and LUMO diagrams of **13** and **14**; blue arrows highlight the lower electron density on the aryl amide in the LUMO. (**d**) Lippert–Mataga plot for **13** and **14**, including calculated μ values. (**e**) Structures of GB_646_-HaloTag ligand (**15**) and HaloTag conjugate (**15–HT**). (**f**) Plot of photon counts/s or particle counts *vs*. time for immobilized **15–HT**; gray shading illumination with 405 nm light. (**g**) SMLM image of a fixed cell expressing histsone H2B– HaloTag and labeled with **15**; the 20,000 frames resulted in 59,165 detected GB_646_ molecules (consolidated emitters); median number of detected photons = 125.2.; scale bar: 5 μm. (**h**) Histogram of localization error; median localization error = 29.5 nm.

To test our hypothesis that excitation of the acylated phthalimide or coumarin lowers nucleophilicity of this amide, we considered model compounds **13** and **14** (**Figure 3b**), which comprise the auxiliary switching groups in photochromic compounds **12** and **5**, respectively. We performed density functional theoretical (DFT) calculations to estimate the change in electron density upon excitation. This *in silico* experiment supports the proposed D–A character of the chromophores as the HOMO shows higher electron density on the amide moieties compared to the LUMO (**Figure 3c**). These experiments suggest that the ICT should decrease the nucleophilicity of the amide upon excitation. To further support this idea, we determined the change in dipole moment upon excitation (Δμ) using a Lippert–Mataga plot (**Figure 3d**).^47, 48^ Plotting the Stokes Shift of the dyes (Δ*v*) *vs*. solvent orientational polarizability (Δf) revealed phthalimide **13** has an estimated Δμ = 3.1 D upon excitation and coumarin **14** exhibits a larger Δμ = 3.6 D. These results suggest that the coumarin lactam system should exhibit sufficient photoinduced ICT to promote the desired photochromic behavior.

We then directly tested the photochromic behavior of the rhodamine–coumarin **5**, which we named “Glitter Bomb 646” (GB_646_). We synthesized the HaloTag^9^ ligand derivative **15** (**Figure 3e**) by reacting known JF_646_-HaloTag ligand with coumarin **4**. We then incubated this compound with purified HaloTag protein to form the conjugate (**15–HT**). Like the parent compound **5**, the HaloTag ligand **15** and the HaloTag conjugate **15–HT** exhibited low p*K*_a_ values, showing no appreciable visible absorption at pH ≥ 2. To evaluate photochromic behavior, the GB_646_-HaloTag protein conjugate (**15– HT**) was deposited on a glass surface and imaged in PBS in the presence of intermittent 405 nm activation light. Increased photon counts and overall number of activated molecules were observed under 405 nm illumination (**Figure 3f**), confirming that incorporating a coumarin into a Si-rhodamine can yield a photochromic derivative. We then tested the utility of **15** in a SMLM experiment. GB_646_-HaloTag ligand (**15**) was incubated with cells expressing the Halo-Tag fused to histone H2B. Illumination with both far-red excitation light (633 nm) and violet activation light (405 nm) allowed imaging of chromatin structure (**Figure 3g**) with reasonable photon output of GB_646_ (*N* = 125.2 photons per localization event) resulting in a localization precision = 29.5 nm (**Figure 3h**).

### A spontaneously blinking dye for super-resolution imaging based on JF_549_

The GB_646_ system confirms our hypothesis that appending a coumarin onto a dye with a relatively low *K*_L–Z_ can produce a photochromic label (**Figure 1b**). We then investigated if a coumarin switching auxiliary could be used to create a spontaneously blinking fluorophore when incorporated into a rhodamine with a higher *K*_L–Z_. We considered compound **8** (**Figure 2a**), which is derived from JF_549_ (**2**; **Figure 1a**); we gave this compound the moniker “Glitter Bomb 549” (GB_549_). Compound **8** exhibits a relatively low p*K*_a_ = 2.9 (**Figure 2c,d**), which could yield a spontaneously blinking dye with a low duty cycle near physiological pH. We were curious whether conjugation to the HaloTag protein could increase this p*K*_a_, as many rhodamine ligands show a shift to the fluorescent, zwitterionic form upon binding their cognate biomolecular target.^12, 49^ To test this hypothesis, we transformed JF_549_-HaloTag ligand (**16**) into the GB_549_-HaloTag ligand **17** through HATU-mediated conjugation with coumarin **4**. We then examined the fluorescence of **17** and **17** bound to purified HaloTag protein (**17–HT**). We observed a marked shift in p*K*_a_ from 2.9 for the free ligand **17** to p*K*_a_ = 4.6 for the HaloTag conjugate; the fluorescence of the dye also substantially increased upon binding (**Figure 4a**). The single molecule behavior of **17–HT** was investigated by sparsely depositing the labeled protein on a glass surface. The spontaneous blinking was clearly visible, with high signal to noise ratio, acceptable duty cycle (0.095%), and acceptable photon yield (*N* = 193 photons/localization event; **Figure 4c**).

**Figure 4.**
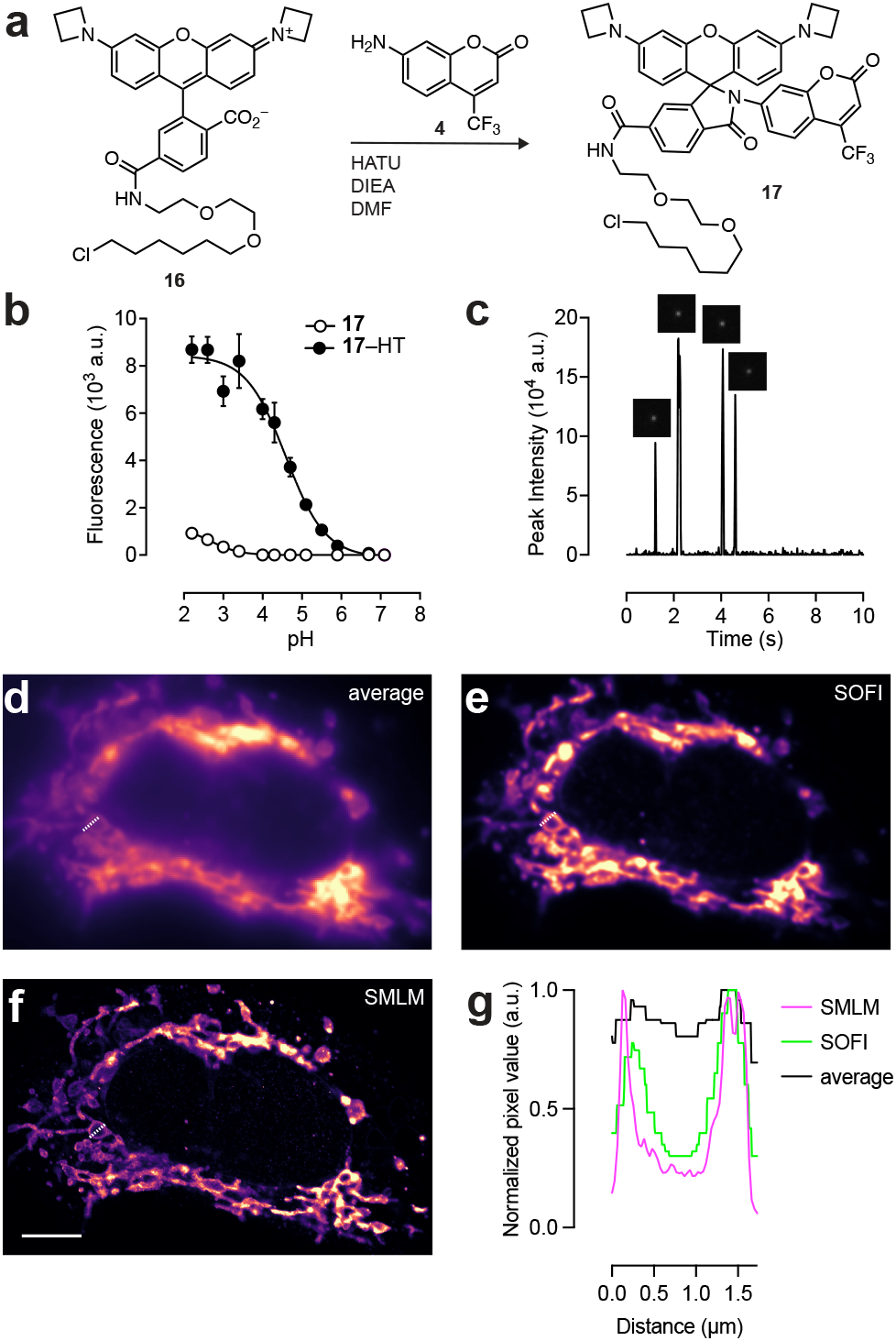
Synthesis and utility of GB_549_. (**a**) Condensation of JF_549_-HaloTag ligand (**16**) and coumarin **4** to synthesize GB_549_-HaloTag ligand **17**. (**b**) Plot of fluorescence intensity *vs*. pH of **17** and **17** bound to HaloTag protein (**17–HT**). Plot of intensity *vs*. time of a single **17–HT** showing four localization events before apparent photobleaching. (**d**–**f**) Imaging experiment of fixed cells expressing TOMM20–HaloTag and labeled with **17**; (**d**) average, (**e**) SOFI, and (**f**) SMLM images; 50,000 frames; 2.3 × 10^6^ localizations; median localization error = 23 nm; scale bar: 5 μm. (**g**) Plot of pixel value *vs*. location for line scan analysis comparing average, SOFI, and SMLM images in panels **d**–**f**.

To evaluate the utility of GB_549_-HaloTag ligand (**17**) in SMLM experiments, we again used cells expressing TOMM20–HaloTag fusion proteins. In a fixed cell preparation, the spontaneously blinking dye showed a satisfactory combination of labeling density and duty cycle with a mean on time of ~20 ms; this allowed assembly of a super-resolution image in <8 min using either super-resolution optical fluctuation imaging (SOFI) or SMLM (**Figure 4d–g**).

### A general utility, spontaneously blinking label based on Alexa Fluor 594

We then considered compound **10**, another candidate spontaneously blinking dye (**Figure 2c,d**) that is based on the optimized and water-soluble fluorophore Alexa Fluor 594. We prepared functionalized derivatives of this “Glitter Bomb 594” (GB_594_) dye starting from the 5-carboxy-AF_594_ methyl ester (**18**; **Scheme 1**). Coupling of **18** with coumarin **4** afforded the rhodamine lactam (**19**), which was then saponified with NaOH to give the 5-carboxy-GB_594_ (**20**). This carboxyl functionality could be further derivatized to prepare derivatives for different attachment strategies including the *N*-hydroxysuccinimidyl (NHS) ester (**21**) for antibody labeling and a phalloidin derivative **22** to directly stain filamentous actin (F-actin).

**Scheme 1.**
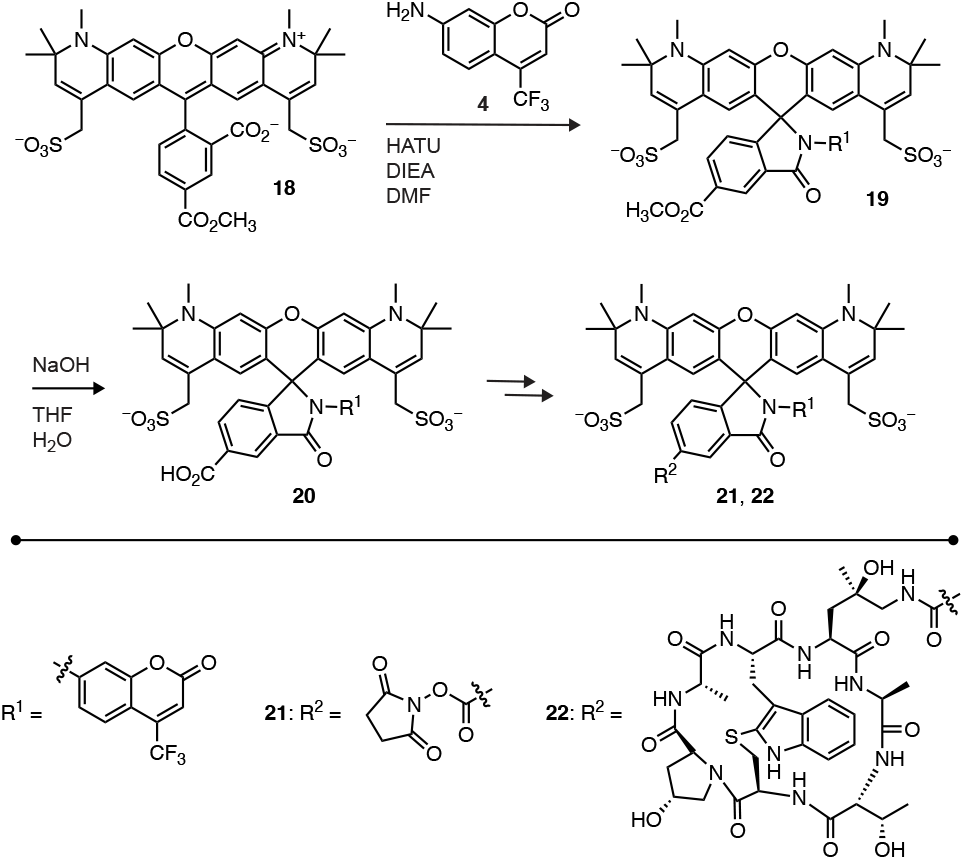
Synthesis of GB_594_ derivatives 21–23.

We first evaluated GB_594_ as an antibody label using the NHS ester **21**. This compound was conjugated to a secondary antibody with a low degree of labeling; dye/antibody < 1. This reagent enabled 3D super-resolution (iPALM)^50^ experiments to image microtubules using a primary anti-tubulin antibody and the GB_594_-labeled secondary (**Figure 5a**). The spontaneously blinking fluorophore exhibited high photon outputs (~1200 photons/localization event) giving high localization precision in all three dimensions (σ ≈ 15 nm). These SMLM images show higher resolution compared to conventional microscopy, showing apparent individual microtubules with full-width at half-maximum (FWHF) measurements ranging from 60–130 nm (**Figure 5a–c**), which is the expected range for anti-body-labeled microtubules.^51^

**Figure 5.**
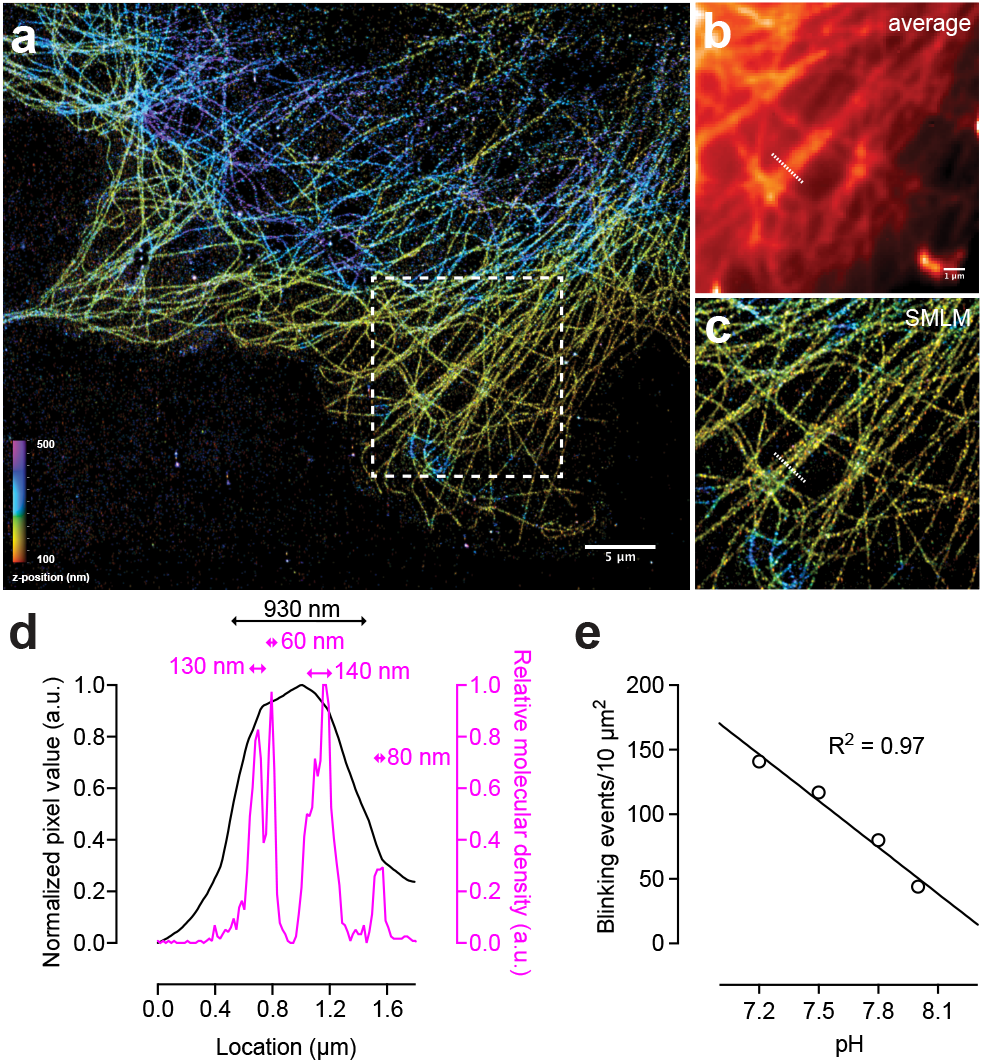
Use of GB_594_-antibody conjugate from 21 for microtubule labeling. (**a–c**) Image of microtubules stained with antibody conjugate prepared from NHS ester **21** with (**a**) 3D super-resolution image of entire field of view (**b**), zoom-in with diffraction limited resolution, and (**c**) zoom-in with super-resolution. (**d**) Plot of pixel value *vs*. location for line scan analysis comparing average and SMLM images in panels **b**–**c**. (**e**) Plot of blinking events per area *vs*. pH for antibody–**21** conjugate; scale bars for all images: 5 μm.

We also used the labeled antibodies to investigate the effect of pH on the blinking properties. The **21**–antibody conjugate was dispersed on a glass slide and the spontaneous blinking behavior of **15** was recorded in buffers with different pH (**Figure 5e**). As expected by this protonation-driven event, more basic buffers reduce the number of localization events in a given unit area over time; the blinking rate decreased by ~60% upon changing pH from 7.2 to 8.0. Adjustment of pH allows additional control over blinking kinetics and could be beneficial when imaging dense structures that require sparser activation to avoid overlapping localizations during the SMLM imaging experiment.

In addition to using an antibody strategy to visualize microtubules, we also imaged F-actin using the small-molecule toxin phalloidin, which binds filamentous actin. This small-molecule label circumvents the linkage error introduced by antibody labeling; the large size of a primary–secondary antibody complex can displace the fluorophore from its biological target by 10–15 nm.^52^ We stained actin with the GB_594_ fluorophore directly using the phalloidin conjugate **22**, which provided a high-resolution and high-quality image (**Figure 6a**) revealing both lamellipodial actin on the leading edge of the cells (**Figure 6b,c**) and actin stress fibers (**Figure 6d,e**).

**Figure 6.**
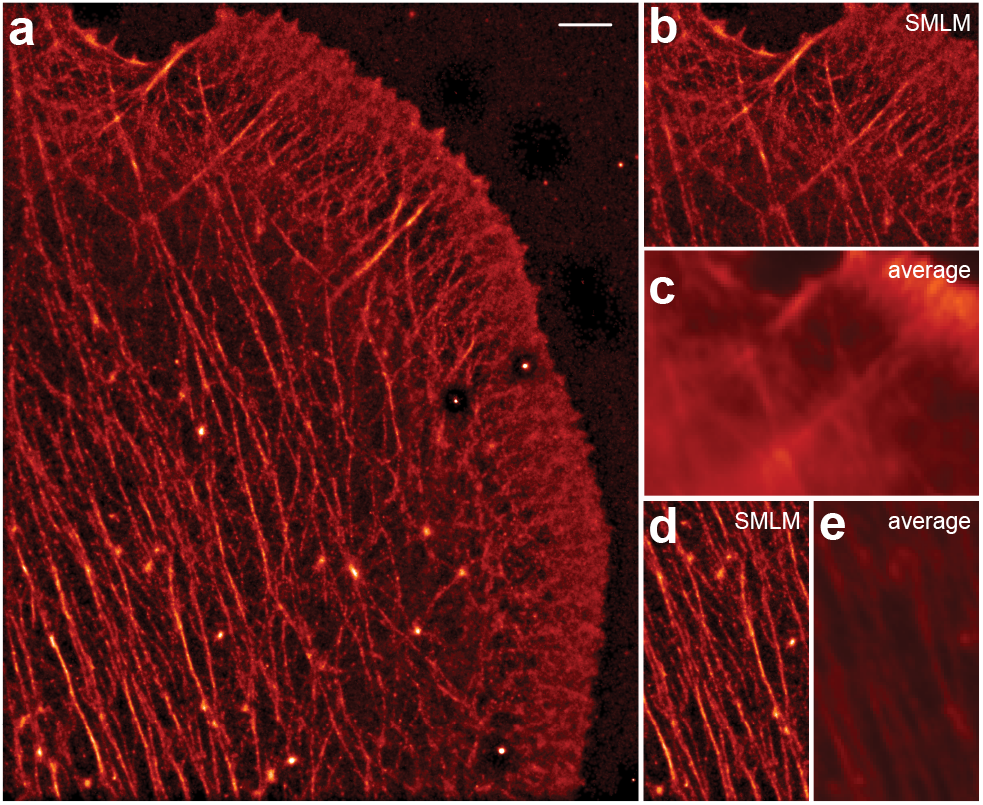
Use of GB_594_-phalloidin (23) for actin labeling. (**a**) SMLM image of the actin cytoskeleton using GB_594_-phalloidin (**23**). (**b**–**c**) Zoom-in of peripheral cellular region showing lamellipodial actin with (**b**) SMLM and (**c**) average renderings. (**d**–**e**) Zoom-in of central cellular region showing actin stress fibers with (**d**) SMLM and (**e**) average renderings. Scale bars for all images: 1 μm.

## CONCLUSION

The last three decades have seen quantal leaps in imaging methods, giving biologists new tools to image cellular components. SMLM is one transformative method that substantially improves the resolution and detail from fluorescence microscopy experiments. This approach relies heavily on both photoactivatable and switchable dyes, making the development of new labels crucial for pushing the frontier of super-resolution imaging. We describe a general strategy to create photochromic and spontaneously blinking dyes using one-step amidation of known rhodamine fluorophores with a classic coumarin laser dye.

This “glitter bomb” approach is based on an understanding of rhodamine and coumarin properties (**Figure 1**,**2**). Rhodamine dyes exist in equilibrium between a lactone and zwitterionic form; the equilibrium constant, *K*_L–Z_, varies widely depending on the chemical structure of the dye.^19^ This complicates the development of spontaneously blinking or photochromic rhodamines since each dye usually requires a specific aryl auxiliary with different properties.^27^ We rationalized that the unique properties of 7-aminocoumarins could overcome this problem. In the ground state, the resonance-driven charge-transfer properties of 7-aminocoumarins result in lower nucleophilicity of the aryl nitrogen. When incorporated into rhodamine dyes with a relatively high *K*_L–Z_, values, the resulting conjugate spontaneously blinks. In the excited state, the coumarin shows an even higher charge-transfer. Attachment of a coumarin to a rhodamine dye with low *K*_L–Z_ yields a photochromic dye.

This combined insight into the basic photophysics of both coumarin and rhodamine dyes generates new labels starting with a farred photochromic fluorophore GB_646_ suitable for cellular SMLM imaging (**Figure 3**). Use of other rhodamine dyes yields spontaneously blinking dyes for fixed-cell experiments including GB_549_ (**Figure 4**) and GB_594_ (**Scheme 1**). In particular, the solubility of GB_594_, which stems from the parent fluorophore Alexa Fluor 594, can be used with standard antibody labeling (**Figure 5**) and phalloidin conjugation (**Figure 6**). We expect exploration of other rhodamine–coumarin combinations to yield additional labels for advanced imaging experiments, leading to a better understanding of the molecular underpinnings of living systems.

## AUTHOR INFORMATION

### Author Contributions

The manuscript was written through contributions of all authors.

### Funding Sources

This work was supported by the Howard Hughes Medical Institute (HHMI). Additional support provided by NIH RO1 GM 117188 (CGG) and NSF 1716316 (JAG).

### Notes

U.S. Patent 11,067,566 describing photochromic and spontaneously blinking xanthene fluorophores (with inventors F.J. and L.D.L.) is assigned to HHMI.

## ACKNOWLEDGMENT

We thank the Janelia Support Teams. This article is subject to HHMI’s Open Access to Publications policy. HHMI lab heads have previously granted a nonexclusive CC BY 4.0 license to the public and a sublicensable license to HHMI in their research articles. Pursuant to those licenses, the author-accepted manuscript of this article can be made freely available under a CC BY 4.0 license immediately upon publication.

## ABBREVIATIONS

SMLM: single-molecule localization microscopy
FP: fluorescent protein
HOMO: highest occupied molecular orbital
LUMO: lowest unoccupied molecular orbital
HATU: hexafluorophosphate azabenzotriazole tetramethyluronium
D–A: donor–acceptor
ICT: intramolecular charge transfer

